# Stable breast cancer prognosis

**DOI:** 10.1101/2021.09.13.460002

**Authors:** Xiaomei Li, Lin Liu, Jiuyong Li, Thuc D. Le

## Abstract

Predicting breast cancer prognosis helps improve the treatment and management of the disease. In the last decades, many prediction models have been developed for breast cancer prognosis based on transcriptomic data. A common assumption made by these models is that the test and training data follow the same distribution. However, in practice, due to the heterogeneity of breast cancer and the different environments (e.g. hospitals) where data are collected, the distribution of the test data may shift from that of the training data. For example, new patients likely have different breast cancer stage distribution from those in the training dataset. Thus these existing methods may not provide stable prediction performance for breast cancer prognosis in situations with the shift of data distribution. In this paper, we present a novel stable prediction method for reliable breast cancer prognosis under data distribution shift. Our model, known as Deep Global Balancing Cox regression (DGBCox), is based on the causal inference theory. In DGBCox, firstly high-dimensional gene expression data is transferred to latent network-based representations by a deep auto-encoder neural network. Then after balancing the latent representations using a proposed causality-based approach, causal latent features are selected for breast cancer prognosis. Causal features have persistent relationships with survival outcomes even under distribution shift across different environments according to the causal inference theory. Therefore, the proposed DGBCox method is robust and stable for breast cancer prognosis. We apply DGBCox to 12 test datasets from different breast cancer studies. The results show that DGBCox outperforms benchmark methods in terms of both prediction accuracy and stability. We also propose a permutation importance algorithm to rank the genes in the DGBCox model. The top 50 ranked genes suggest that the cell cycle and the organelle organisation could be the most relevant biological processes for stable breast cancer prognosis.

**Author summary:** Various prediction models have been proposed for breast cancer prognosis. The prediction models usually train on a dataset and predict the survival outcomes of patients in new test datasets. The majority of these models share a common assumption that the test and training data follow the same distribution. However, as breast cancer is a heterogeneous disease, the assumption may be violated in practice. In this study, we propose a novel method for reliable breast cancer prognosis when the test data distribution shifts from that of the training data. The proposed model has been trained on one dataset and applied to twelve test datasets from different breast cancer studies. In comparison with the benchmark methods in breast cancer prognosis, our model shows better prediction accuracy and stability. The top 50 important genes in our model provide clues to the relationship between several biological mechanisms and clinical outcomes of breast cancer. Our proposed method in breast cancer can potentially be adapted to apply to other cancer types.

## Introduction

Breast cancer prognosis can help tailor treatments for patients and improve their survival outcomes. Traditional breast cancer prognosis mostly relies on the tumour-node-metastasis (TNM) staging system [1]. However, tumours with the same clinical characteristics can differ in both prognosis and treatment response because breast cancer is heterogeneous with variable molecular mechanisms of carcinogenesis and cancer development. Therefore, it is desirable to use transcriptomic data to conduct breast cancer prognosis at molecular level.

In the past decades, a large amount of transcriptomic data have been generated for breast cancer using next-generation sequencing and microarray technologies, and many computational methods have been developed to utilize the data for breast cancer prognosis. Most of the methods are based on Cox regression [2]. For example, rorS [3] trains a risk model by ridge penalized Cox regression based on the breast cancer subtype information obtained from gene expression data. TAMR13 [4] builds a Cox regression model based on the data of gene probe cluster centroids. Similarly, OncotypeDX [5], GENIUS [6] and EndoPredict [7] use Cox regression to determine the weights of the selected genes and then calculate the hazard risk as a weighted combination of the gene expression levels.

Some machine learning based methods have also been used for cancer prognosis, such as Cox regression regularized by elastic Net [8], Random Survival Forest (RSF) [9], Support Vector Regression for Censored Data (SVRc) [10], Cox proportional hazards deep neural network (DeepSurv) [11] and so on (see [12] for a review). These methods take advantage of machine learning techniques, to selecting vital features in high-dimension transcriptomic data and to model non-linear relationships between features (genes) and survival outcomes.

These methods assume no distribution shift from the training data to the test data (new unseen data). However, the assumption will be violated in real applications. For example, gene expression data may be produced using different platforms, by different labs and patients may come from different countries or have different breast cancer stages. Users do not know what future unseen data will be like. Inaccurate prognosis predictions will have resulted from the distribution shift even though the models perform well in the training and hold out validation datasets.

The computational methods need to have stable prediction performance since their predictions directly impact treatment decisions. Here the stability of a method means that the method performs consistently with the training data and the test data (i.e. unseen breast cancer patients) which may have different distributions. One strategy to improve the stability of a prediction method is to utilize samples from multiple sources to train the model. For example, several methods have been developed to select a set of robust features from multiple datasets obtained from the studies of the same disease, and use these features to build a more reliable prediction model [13, 14]. Another strategy is to employ transfer learning techniques to train a robust prediction model [15]. For example, the method in [16] transfers the weights of a variational auto-encoder trained with all the transcriptomic data of 20 cancer types (source domain) to the survival prediction model for patients from a single cancer type (target domain). The transfer learning based methods enhance the robustness of survival predictions in unseen cancer patients by extracting the shared genomic knowledge from the source domain and transfer the knowledge to the target domain.

From the above discussion, we can see that these existing stable prediction methods, when used for breast cancer prognosis, require cancer patient samples with transcriptomic data and/or survival outcomes coming from either different sources or multiple domains in the training phases. Both are challenging in practice because it is time-consuming to follow up cancer patients over long-term periods and collect a wide variety of data. Therefore, it is useful to develop survival prediction models that train on an available dataset and are robust to distribution shifts in test data from new environments.

In this paper, we focus on designing a model for stable survival prediction under new environments where the data distribution has been shifted from that in the training dataset. The new environment is agnostic to us and the distribution in the test data is unknown. In this case, we cannot use any information from the test data when we train a model. To learn a robust prediction model across different environments, the predictors (signatures) must be causal genes, i.e. genes causing the development of the disease [17]. The pinpointed causal genes in the model can be translated into disease mechanism and clinical treatment [18]. Therefore, we will model stable breast cancer prognosis predictions as a causal problem.

In the research area of causal inference, the causal effects are used for measuring the impact of a treatment on patients. Covariate balancing strategies are used for estimating the causal effect of a treatment variable on the outcome in the observational data. A recent global balancing strategy by Kuang *et al*. [19] is a covariate balancing strategy used to estimate the causal effects of covariates on the outcome when one does not know which covariates are causal and which ones are not. This can be used for our design because we do not know which genes are causal. However, three limitations prevent this method from being immediately applied to achieve a stable breast cancer prognosis. Firstly, the method is for a classification problem with the categorical outcome not for a regression problem with the continuous outcome as in prognosis prediction. Secondly, this method is only applicable to data without censoring, unfortunately, the survival outcomes in breast cancer datasets are almost always censored. Thirdly, it cannot deal with high dimensional gene expression data used in building breast cancer prognosis models.

We propose a novel method, Deep Global Balancing Cox regression (DGBCox) for stable breast cancer prognosis based on censored survival data and gene expression data. With DGBCox, we have made two main contributions: (1) We leverage the strengths of both regularized Cox regression and global balancing approach and propose a novel method for stable cancer prognosis. To the best of our knowledge, this is the first cancer prognosis model which is aimed at stable predictions across different environments. (2) We have developed a new deep learning based framework to deal with high dimensional gene expression data.

We apply DGBCox to breast cancer datasets from different studies. The experiment results have demonstrated that DGBCox performed better than baseline methods and existing breast cancer prognosis methods with regard to accurate and stable breast cancer prognosis. We interpret the important genes for the stable breast cancer prognosis in our model. A high percentage of the top 50 important genes discovered by DGBCox has been selected by previous research for breast cancer prognosis. The important genes are enriched in several well-known biological processes involved in the development of cancer. These findings may help to understand the biological mechanisms of breast cancer development and progression.

## Materials and methods

### Datasets

In this study, we use 13 genome-wide expression datasets containing in total 5668 breast cancer patients from different repositories (Table 1). Among these datasets, TCGA753 and TCGA500 are subsets of TCGA1093 generated by The Cancer Genome Atlas (TCGA) program (https://www.cancer.gov/tcga). METABRIC is available at the European Genome-phenome Archive (EGA) (https://www.ebi.ac.uk/ega/, accession number EGAS00000000083, access required). The MAINZ, TRANSBIG, UPP, UNT, and NKI datasets are from Bioconductor (https://bioconductor.org/) data packages, breastCancerMAINZ, breastCancerTRANSBIG, breastCancerUPP, breastCancerUNT, and breastCancerNKI, respectively. The remaining datasets are all downloaded from the Gene Expression Omnibus database (https://www.ncbi.nlm.nih.gov/geo/). More details about the datasets are in Section 1 in S1 File.

**Table 1.** A summary of the datasets.

| Dataset | Platform | #Samples | Min time<sup>1</sup> | Max time<sup>2</sup> | Event (%)<sup>3</sup> | Reference |
| --- | --- | --- | --- | --- | --- | --- |
| METABRIC | Illumina HumanHT-12 V3 | 1283 | 0 | 29.60 | 32.68 | [20] |
| TRANSBIG | Affymetrix Human Genome U133A | 198 | 0.34 | 24.95 | 31.31 | [21] |
| UNT | Affymetrix Human Genome U133A/B | 137 | 0.17 | 14.53 | 21.05 | [22] |
| UPP | Affymetrix Human Genome U133A/B | 251 | 0.08 | 12.75 | 23.08 | [23] |
| MAINZ | Affymetrix Human Genome U133A | 200 | 0.08 | 19.72 | 23.00 | [24] |
| NKI | Agilent Human Genome Microarray | 337 | 0.02 | 18.35 | 34.17 | [25, 26] |
| GSE6532 | Affymetrix Human Genome U133 Plus 2 | 414 | 0.02 | 16.85 | 34.66 | [27] |
| GEO736 | Affymetrix Human Genome U133 Plus 2 | 736 | 0 | 18.52 | 47.42 | [4, 28–31] |
| TCGA1093 | Illumina GA/HiSeq RNA-seq | 1093 | 0 | 23.44 | 10.35 | [32] |
| TCGA753 | Illumina GA/HiSeq RNA-seq | 753 | 0 | 17.90 | 9.57 | [32] |
| TCGA500 | Illumina GA/HiSeq RNA-seq | 500 | 0.003 | 17.90 | 9.02 | [32] |
| UK | Illumina humanRef-8 V1 | 207 | 0.39 | 10.00 | 37.20 | [33] |
| HEL | Illumina HumanHT-12 V3 | 115 | 0 | 5.00 | 21.74 | [34] |

1: the minimum survival time in years in a dataset. 2: the maximum survival time in years in a dataset. 3: the percent of patients experiencing events in a dataset.

The clinical outcomes included in the datasets used in this study include relapse free survival time (UPP, GSE6532, GEO736, TCGA1093, TCGA753, TCGA500, METABRIC, and UK), distant metastasis free survival time (TRANSBIG, UNT, MAINZ, and NKI), and death or distant metastasis free survival time (HEL). In all the datasets, the unit of survival time is a year. The maximum survival time in the datasets ranges from 5 years to 29.6 years. The medical event in the datasets is breast cancer relapse, metastasis or death. The event percentage in the datasets ranges from 9.02% to 47.42%. These breast cancer datasets were collected from different batches of experiments with different RNA-sequencing platforms and different pipelines for RNA-seq data processing and have different ranges of survival time and different ratios of event occurrences. Therefore, it is reasonable to assume that the distributions of these datasets are different.

### Stable cancer prognosis using Deep Global Balancing Cox regression (DGBCox)

#### Overview of DGBCox

As illustrated in Fig 1, DGBCox has three main components in the training phase, a deep auto-encoder neural network (DAE), a survival global balancing component (SGB) and a regularized Cox regression (Cox). DAE is used for extracting the latent patient representations from the original gene expression data of a patient. Cox is used to predicting a patient’s hazard risk that indicates how likely the patient is to experience an event. SGB re-weights the training data samples so that it is possible to infer the causal relationships between latent features and the survival outcomes. The three components of DGBCox are trained using the input dataset *D* = (**X**, *T, E*), where **X** is gene expression data, *T* is observed survival time and *E* is event status. In the test phase, DGBCox predicts the hazard risk of a patient only based on the trained DAE and Cox components. In the following sections, we explain the three components, the training and test phases of DGBCox.

**Fig 1.**
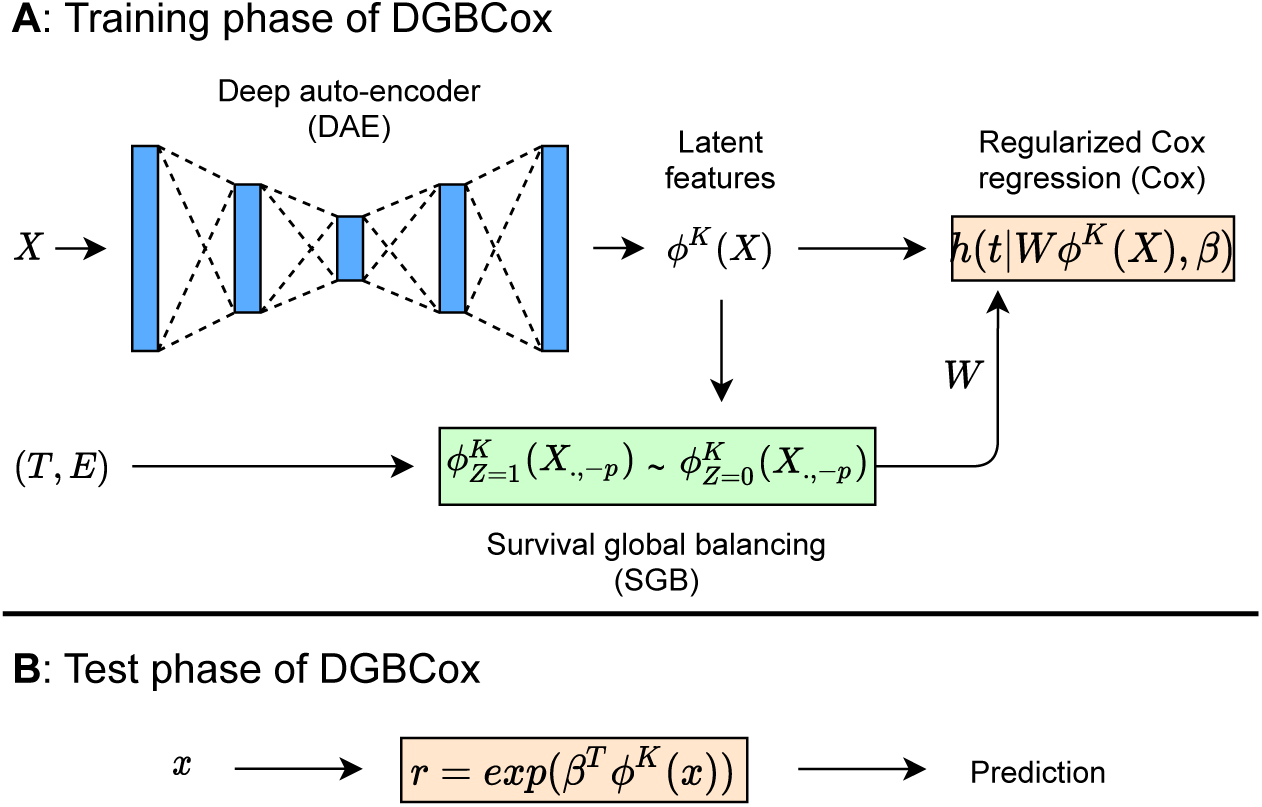
The framework of DGBCox. **A**. DGBCox contains three main components in the training phase, i.e. DAE, SGB and Cox. SGB balances weighted covariate distributions in treated and control groups and then offers weights in *W* to Cox. DAE provides latent representations in *ϕ*^*K*^(**X**) to SGB and Cox. DAE, SGB and Cox are jointly trained by an optimization algorithm. **B**. For an unseen test patient with gene expression profile *x*, the survival prediction is an exponent of the linear combination of latent features, where the coefficients *β* (in Cox) and the encoder function *ϕ*^*K*^(·) (in DAE) are determined in the training phase.

#### Deep auto-encoder neural network (DAE)

The gene expression data of cancer patients are typical of high dimensionality, but the global balancing technique employed by DGBCox (for the SGB component) cannot deal with high dimensional data if it successively treats each gene as a treatment variable and learns weights to balance the dataset (details in the next section). Therefore we apply a deep auto-encoder neural network [35] to reduce the dimensionality of the gene expression data and feed the low dimensional latent features to SGB.

DAE consists of two parts, the encoder and the decoder, which can be defined as transitions *ϕ* : **X** → *ϕ*(**X**) and *φ* : *ϕ*(**X**) → *φ*(**X**), respectively. Specifically, the latent representations for each hidden layer of the encoder can be given as:

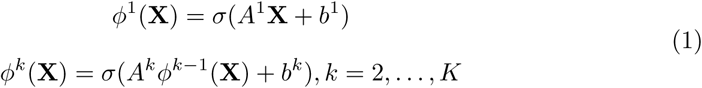

where *K* is the number of hidden layers in the encoder. *A*^*k*^ and *b*^*k*^ are the weight matrix and bias vector on the *k*-th hidden layer of the encoder, respectively. *σ*(.) is an activation function.

The decoder transitions the resulting latent representations *ϕ*^*K*^(**X**) to a reconstructed 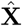 with the same shape of **X**:

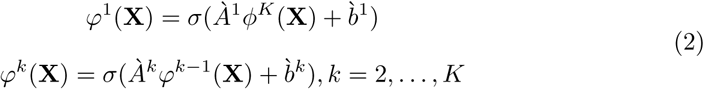

where *À*^*k*^ and 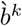 are the weight matrix and bias vector on the *k*-th layer of the encoder respectively. The reconstructed 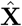 is the *K*-th layer output *φ*^*K*^(**X**). DAE is trained to minimize reconstruction errors between the input gene expression data **X** and the reconstructed 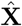, such as 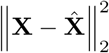.

The output of the DAE component is the low-dimensional latent features (in *ϕ*^*K*^(**X**)) which are the non-linear combinations of the input gene expression data. For the sake of simplicity, the superscript *K* is dropped when the context is clear. *ϕ*(**X**) is used for input data of SGB and Cox instead of **X** as described in the following sections.

#### Survival global balancing (SGB)

It is known that causal relationships between covariates are persistent and stable across different environments whereas noncausal relationships or purely correlations may variant across different environments [36]. Therefore to achieve stable predictions for cancer prognosis, for our proposed DGBCox, we aim to use the latent features which are causally related to the survival outcomes (called causal latent features in this paper) as the predictors in the prediction model (i.e. the Cox regression model) for prognosis. To find the causal features, we need to estimate the causal effect of a latent feature on the survival outcomes. Covariate balancing is a way to estimate causal effects [37]. Here we extend the global balancing method [19] to censored survival data and propose the survival global balancing (SGB) component for DGBCox. SGB re-weights the training data samples, which enables estimation of the causal effect of each latent feature and thus to identify the causal features.

In the following, we start with introducing the general idea of covariate balancing and the global balancing method [19] for non-censored data. Then we propose our method to extend the covariate balancing approach and the global balancing method to censored survival data. Lastly, we incorporate the extended global balancing approach with DAE for censored data.

Let *Z*_*i*_ ∈ {0, 1} denote treatment assignment, with *Z*_*i*_ = 1 indicating the *i*-th patient is treated and *Z*_*i*_ = 0 indicating the opposite. **X**_*i*_ and *Y*_*i*_ describe the features and the observed survival time for the *i*-th patient respectively. The subscript *i* will be dropped when the context is clear. *Y* ^(*z*)^ (*z* ∈ {0, 1}) denote the potential survival outcomes of a patient under the treatment assignment *Z* and *Y* = *ZY* ^(1)^ + (1 − *Z*)*Y* ^(0)^. The observed dataset is composed of i.i.d. samples (*Y*_*i*_, *Z*_*i*_, **X**_*i*_), for *i* = 1, 2, …, *N*, where *N* is the sample size of the training dataset. In causal inference, to estimate of average treatment effect from observed data with non-random treatment assignment, covariate balancing is a commonly used method to balance the distribution of the covariates between treated and control groups [19]. Under the assumption of unconfoundedness (*Z* ╨ (*Y* ^(1)^, *Y* ^(0)^)|**X**), the general idea of the covariate balancing approach is to re-weight the training samples, i.e. re-weigtht the *i*-th sample with the weight *W*_*i*_, and the weight vector *W* = {*W*_1_, *W*_2_, …, *W*_*N*_} is obtained as follows:

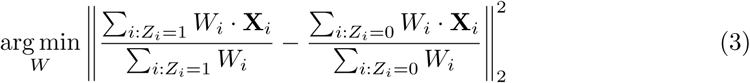

Through the sample re-weighting, the average treatment effect can be estimated using the difference of the expected weighted survival time of patients in treated and control groups [19].

When the causal features (regarded as treatments) are unknown, Kuang *et al*. proposed a Global Balancing (GB) method to identify causal features from the observational data [19]. GB successively takes each covariate as a treatment variable and balance the distribution of all the other covariates between treatment and control. As a result, GB learns global sample weights in *W* satisfied with

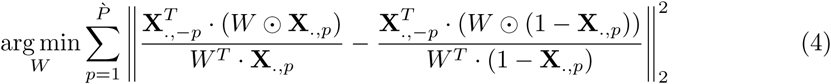

where 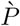 is the dimension of **X, X**_.,*p*_ is the *p*-th covariate or the binary of the *p*-th covariate (for non-binary variable) in **X**, and **X**_.,−*p*_ = **X** \ **X**_.,*p*_ holds all the remaining covariates except the *p*-th covariate in **X**. ⊙ refers to element-wise multiplication.

Now we extend Formula 3 to right censored survival data. Right censoring occurs when a patient does not experience an event but she is lost to follow-up or the medical study ends. Therefore, when right censoring occurs to a patient, instead of observing *Y*, one observes *T* = *ZT* ^(1)^ + (1 − *Z*)*T* ^(0)^, and *T* ^(*z*)^ = *min*(*Y* ^(*z*)^, *C*), where *C* denotes the censoring time. Similarly, one observes the event status *E* = *ZE*^(1)^ + (1 − *Z*)*E*^(0)^, where *E*^(*z*)^ = *I*(*C* ≥ *Y* ^(*z*)^) and *I*(·) is an indication function. *E*_*i*_ = 1 indicates an event has been observed for the *i*-th patient and the realized survival time of the patient has been completely observed, while *E*_*i*_ = 0 indicating the opposite. The censored survival data are thus samples of i.i.d. random vectors (*T, E, E* · *Y, Z*, **X**). Similar to data without censoring, we assume the treatment variable is independent of censored and uncensored survival time given the gene expression data, the unconfoundedness assumption is now extended to (*T* ^(1)^, *T* ^(0)^, *Y* ^(1)^, *Y* ^(0)^) ╨ *Z*|**X**. In addition, we assume the censoring is independent of survival time and gene expression data given the treatment variable (*T* ^(1)^, *T* ^(0)^, *Y* ^(1)^, *Y* ^(0)^, **X**) ╨ *C*|*Z*). Inspired by estimating of average treatment effect for censored survival data [38], we extend the covariate balancing approach to censored survival data by:

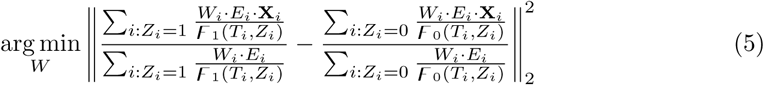

where *F*_*z*_(*T, Z*) = *Pr*(*C > T* |*Z* = *z*) denotes the conditional censoring distribution which is estimated by the Kaplan–Meier method [39, 40].

With the extended covariate balancing approach, we propose SGB to learn global sample weight vector *W* from censored survival data:

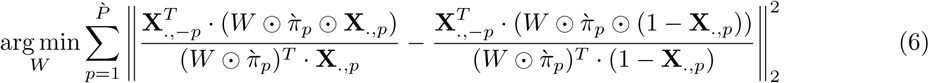

where 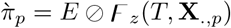, and ⊘ refers to element-wise division.

In gene expression data **X**, the dimension 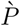 is large. There may be no sufficient samples and computational resources to estimate *W* by optimizing Formula 6. Therefore we successively regard each latent feature in a low-dimensional space from DAE as a treatment variable and learn *W* by:

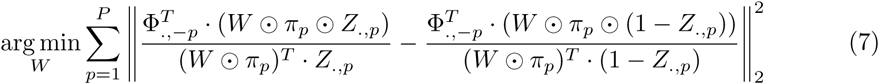

where Φ = *ϕ*(**X**), *P* is the dimension of Φ, and *Z*_.,*p*_ is the *p*-th treatment variable via dichotomizing Φ_.,*p*_ by its mean value. Φ_.,*p*_ is the *p*-th covariate in Φ, and Φ_.,−*p*_ = Φ \ Φ_.,*p*_. π_*p*_ = *E* ⊘ *F* _*z*_(*T, Z*_.,*p*_).

#### Regularized Cox regression (Cox)

In survival analysis literature, a hazard function is used for modelling the instant probability of an individual will experience an event at time *t* given that she has already survived up to time *t*. One can estimate the hazard risk score(s) using the hazard function for each individual based on their baseline data. Cox regression [2] is a commonly used survival analysis method. Here we first introduce the general Cox regression model build on original gene expression data. The hazard function of Cox regression is defined as:

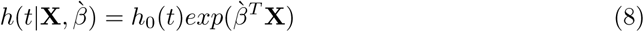

where *h*_0_(*t*) is the baseline hazard function which is not specified in Cox regression. The prediction of Cox regression is the ratio 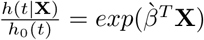 which represents the relative hazard risk of a patient. 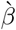 holds the coefficients of the predictors and can be estimated from training data, and this is often done through minimizing the following loss function (called average negative log partial likelihood):

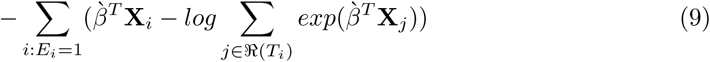

where ℜ(*T*_*i*_) = {*j* : *T*_*j*_ ≥ *T*_*i*_} is the set of patients who are still at risk of event at time *T*_*i*_.

However, Cox regression for high-dimensional data is challenging and often does not behave well [8]. Thus we build a regularized Cox regression model based on the latent features found by DAE instead of the original gene expression data. Moreover, to estimate the causal effects of latent features on survival outcomes, we need to re-weighting training samples by the weights obtained by SGB (Formula 7). Therefore, we use the following loss function of our modified Cox regression:

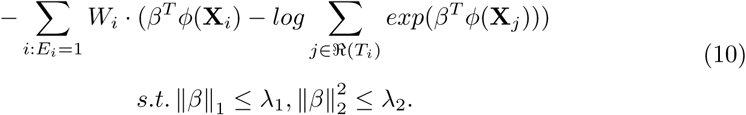

where the L1 regularization (also known as lasso regularization) term (∥*β*∥_1_ ≤ *λ*_1_) and the L2 regularization term 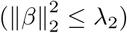 help to avoid overfitting of our modified Cox regression model. After survival global balancing, causal latent features are more likely selected as predictors, which leads to stable performance for cancer prognosis.

#### Joint optimization of the three components in DGBCox

A joint optimization strategy is used for tradeoffs between the prediction accuracy (by Cox), the reconstruction error (by DAE) and the global balancing regularization (by SGB) in DGBCox. The loss function of DGBCox is defined as:

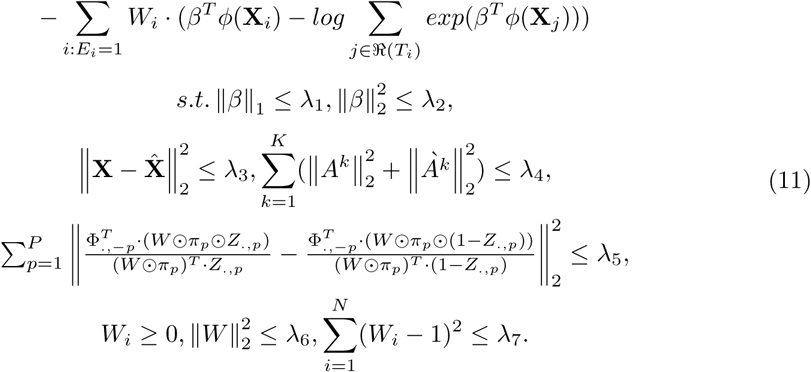

The condition of 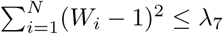 helps to avoid all the weights to be 0.

#### Cancer prognosis with DGBCox

The relative hazard risk score for a new patient is calculated by *r* = *exp*(*β*^*T*^ *ϕ*(*x*)). The patient with a low (hazard) risk score is expected to have a better prognosis than the one with a high risk score.

### Evaluation metrics

We first introduce the traditional evaluation metrics for cancer prognosis methods, and then define the metrics for evaluating the stability of a prognosis method.

#### Concordance index

C-index [41] is a commonly used measure for evaluating the accuracy of the risk score prediction of a cancer prognosis method. Let *Y*_*i*_ and *r*_*i*_ be the potential survival time and the risk score for a patient *i*, respectively. C-index is defined as the concordance probability *Pr*(*r*_*j*_ *> r*_*i*_|*Y*_*j*_ *< Y*_*i*_) for all randomly selected pairs of patients (*i, j*). The concordance means that the patient with a high risk score survives shorter than the one with a low risk score. In other words, the predicted risk score should be negatively associated with the length of survival. However, the potential survival time *Y* cannot be observed in the right censored survival data. The observed survival outcomes contain observed survival time *T* and event status *E*. For the right censored survival data, the concordance probability is extended to *Pr*(*r*_*j*_ *> r*_*i*_|*T*_*j*_ *< T*_*i*_, *T*_*i*_ ≤ *C*_*i*_), where *C*_*i*_ is the right censoring time. *T*_*i*_ ≤ *C*_*i*_ indicates that the patient *i* experienced an event during the follow-up time and *E*_*i*_ = 1. Therefore, C-index can be computed with the following formula:

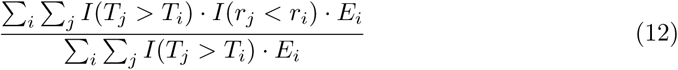

C-index ranges from 0 to 1. If the C-index of a method equals 0.5, that means this method is no better than a random guess model. A model with a C-index of 1 indicates that the model’s predictions and survival outcomes are perfectly concordant. On the contrary, a model with a C-index of 0 means that the model’s predictions and survival outcomes are discordant. In this case, the predicted risk score is positively associated with the length of survival which is not true in cancer prognosis.

#### Hazard ratio

A commonly used metric for evaluating risk group predictions is hazard ratio [42]. We obtain the predicted groups *G* for patients via dichotomizing the predicted hazard risk scores by their median value. If a patient’s risk score is bigger than the median predicted hazard risk score of the cohort, the patient is put into the high-risk group, otherwise, the patient is put into the low-risk group. Then the risk groups *G* is fitted in a Cox regression model:

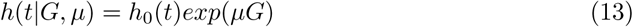

where *h*_0_(*t*) is the same as that in Eq 8. The quantity *exp*(*µ*) is defined as hazard ratio which indicates the risk difference between the two groups of patients. The method with the high hazard ratio performs better than the one with the low hazard ratio.

#### The Log-rank test

We use the Log-rank test [43] to assess whether a method successfully stratified patients into two risk groups with statistically significant different survival distributions. In the Log-rank test, the null hypothesis is that there is no difference between the survival distribution in the two risk groups at any time point. If the p-value of the Log-rank test for a method is less than 0.05, we reject the null hypothesis and consider that the method successfully stratified patients into two risk groups with distinct survival patterns.

#### Metrics for stability evaluation

Given a set of test datasets (*D*^1^, *D*^2^, …, *D*^*M*^), *CI*^*m*^ and *HR*^*m*^ are the C-index and hazard ratio on dataset *D*^*m*^, *m* ∈ {1, 2, …, *M*}, where *M* is the number of test datasets. To assess whether a cancer prognosis model is a stable and accurate model, we define a set of stability evaluation metrics as below.

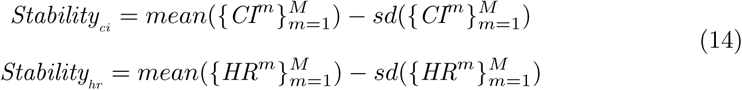

where *mean*(·) and *sd*(·) returns the mean and standard deviation of a given vector that holds C-indices or hazard ratios of a method on all the *M* test datasets. The larger *Stability*_*ci*_ or *Stability*_*hr*_ is, the more stable the cancer prognosis method will be. In this study, a cancer prognosis model with high stability implies that the model has not only a high average accuracy but also a low standard deviation of accuracy across different environments.

### Prioritizing stable features

To interpret the DGBCox method, we propose a permutation importance approach to measure how much DGBCox’s stability depends on the information in each covariate (gene). The permutation importance algorithm calculates the decrease in prediction stability when a covariate is permuted (randomly shuffled). The pseudocode of the permutation importance algorithm is illustrated in Algorithm 1 in S1 File.

### Experimental setting

To evaluate the effectiveness of DGBCox, we create and implement the following baselines to investigate the contribution of the components (in particular DAE and SGB) proposed for DGBCox.

- A Cox regression model with elastic net regularization (Coxnet)
- Deep Auto-Encoder Cox regression (DAECox)

The first baseline (Coxnet) is to evaluate the effect of both DAE and SGB, and the second baseline (DAECox) is to evaluate the effect of SGB. Only integrating SGB and a regularized Cox regression is impractical to the high dimensional gene expression data, therefore, we do not compare DGBCox with such a method. The details of the two baselines are described in Section 2 in S1 File.

Since the proposed method aims at stable prediction of the hazard risks of new breast cancer patients from unknown environments, we train the proposed method and baselines based on METABRIC and test the models on the other 12 independent datasets. C-index is the primary metric to evaluate the accuracy for risk score prediction of a cancer prognosis method. Thus, in this study, the hyper-parameters of the methods are determined by maximizing *Stability*_*ci*_ on validation datasets by a Bayesian hyper-parameters search [44]. More information about the hyper-parameters and the training data partition is in Section 3 in S1 File.

We also compare our methods with the state-of-the-art methods, including DeepSurv [11], Robust [13], AURKA [45], ESR1 [45], ERBB2 [45], GGI [22], GENIUS [6], Endopredict [7], OncotypeDx [5], TAMR13 [4], PIK3CAGS [27], GENE70 [25] and rorS [3]. We implement DeepSurv and Robust as follows. The optimal hyper-parameters of DeepSurv are selected based on cross validation on METABRIC by the Bayesian hyper-parameters Optimization. Robust is trained based on METABRIC to determine the parameters of its predictive model. The remaining methods have selected gene signatures (predictors) based on prior knowledge and clinical information. These methods have been implemented by R package CancerSubtypesPrognosis (https://github.com/XiaomeiLi1/CancerSubtypesPrognosis).

## Results

### Effectiveness of DGBCox

Fig 2A shows the risk score prediction performance of DGBCox and the two baselines (Coxnet and DAECox). Fig 2B shows the risk group prediction performance of the three methods. Fig 2 also shows the stability results based on *Stability*_*ci*_ and *Stability*_*hr*_. Please note that the higher the better for all the evaluation metrics.

**Fig 2.**
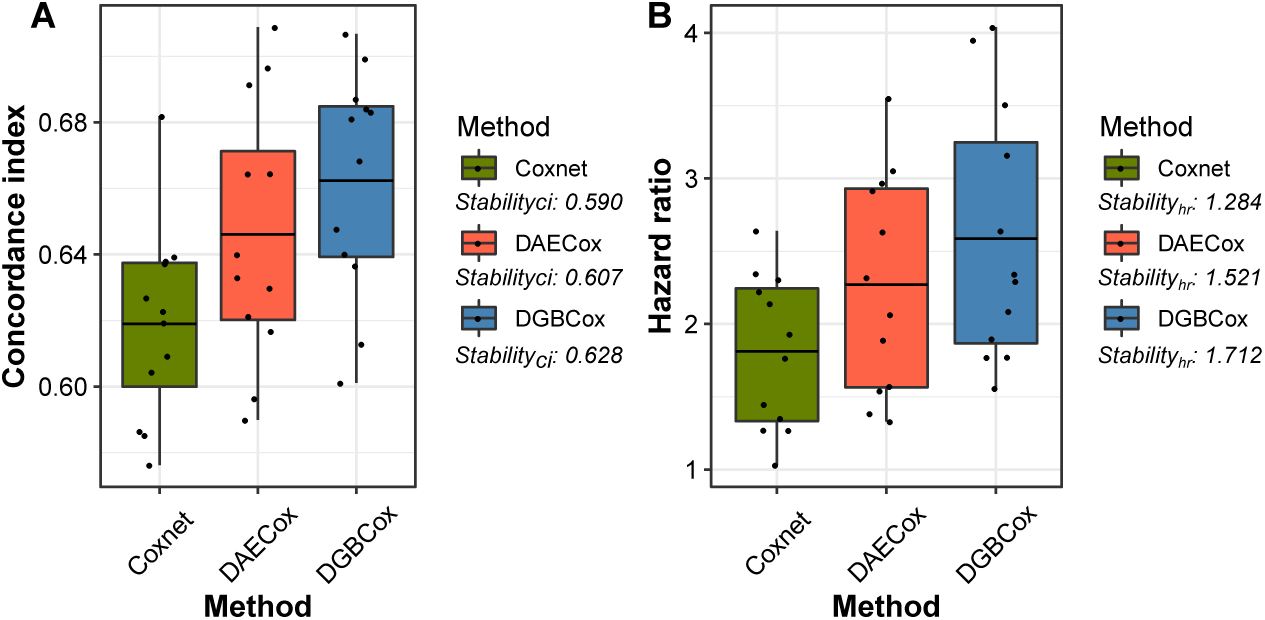
The box plots for the performance of DGBCox and baselines on the independent test datasets. **A:** the box plot for the concordance indices. **B:** the box plot for the hazard ratios. The horizontal line in each box indicates the mean value.

From the results shown in Fig 2, we have the following observations:

- Compared with Coxnet, DAECox achieves better performance in terms of *Stability*_*ci*_ and *Stability*_*hr*_. This indicates that the DAE component improves the performance of Coxnet for both risk score and risk group predictions. This is because Coxnet only models the linear relationship between genes and survival outcomes, while DAECox can capture the non-linear relationship between genes and survival outcomes.
- In comparison with DAECox and Coxnet, DGBCox makes the most stable risk score and risk group prediction. This is because the SGB component ensures approximate estimation of the causal effect of features, therefore, causal features can be selected in the prediction model, which leads to accurate and stable prediction across different environments. Moreover, DGBCox obtains the highest average C-index of 0.662 and the highest average *HR* of 2.586, outperforming the other two methods.

Furthermore, we use the Log-rank test to analyze whether the survival pattern in one risk group is significantly different from that in another one. The results are shown in Fig 3. From the figure, we see that DGBCox outperforms Coxnet and DAECox by 5 and 3 respectively in terms of the number of datasets where a method obtains small p-values (less than 0.05). The results confirm that DGBCox is a viable approach for improving the performance in stratifying breast cancer patients into two risk groups that have distinct survival patterns.

**Fig 3.**
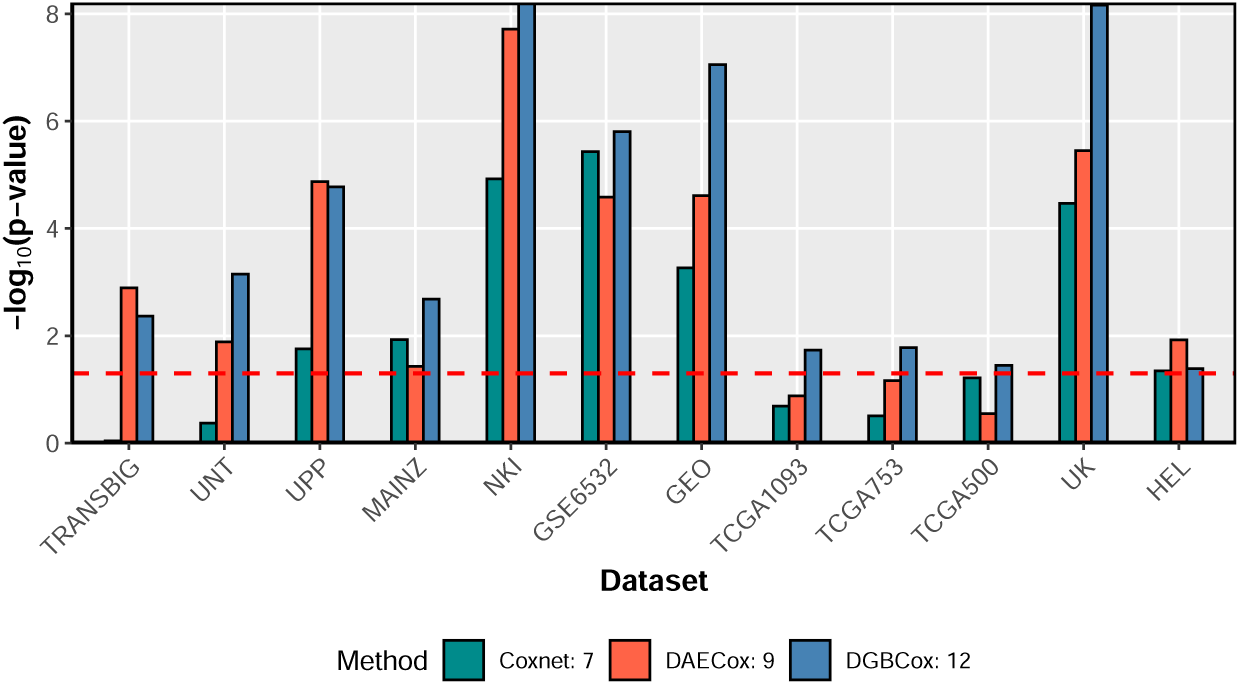
The bar plot for the performance of DGBCox and baselines on 12 test datasets. The x-axis stands for the dataset, and the y-axis shows the log transformation of the p-value of the Log-rank test. The red horizontal line indicates the threshold for significantly results i.e. the results over the red line are statistically significant, while the results under the line are not.

### Comparison with the state-of-the-art methods

Fig 4 shows the C-indices of DGBCox and each of the 13 state-of-the-art methods compared on each of the 12 test datasets, and the *Stability*_*ci*_ of each method across all test datasets. As shown in Fig 4, DGBCox outperforms the state-of-the-art methods on five datasets (UNT, UPP, GSE6532, TCGA1093, TCGA500) in terms of C-index. Meanwhile, the state-of-the-art methods only display the best results on up to two datasets. For a model which is capable of making predictions correctly, the C-index of the model on each test dataset should be higher than 0.5. From Fig 4, we can see that most methods are capable of making predictions correctly on most test datasets, but DGBCox achieves the most notable performance with the C-indices over 0.6 on all the datasets. We observed that the C-indices of some methods (ESR1, ERBB2, Robust, and PIK3CAGS) are less than 0.5 on many datasets, which means the correlations between the predicted risk scores by those methods and the survival outcomes of patients in different cohorts are contradictory. These incoherent associations may be attributable to that some signatures used in these methods vary across different environments, while the Cox regression method cannot correct the distribution shift.

**Fig 4.**
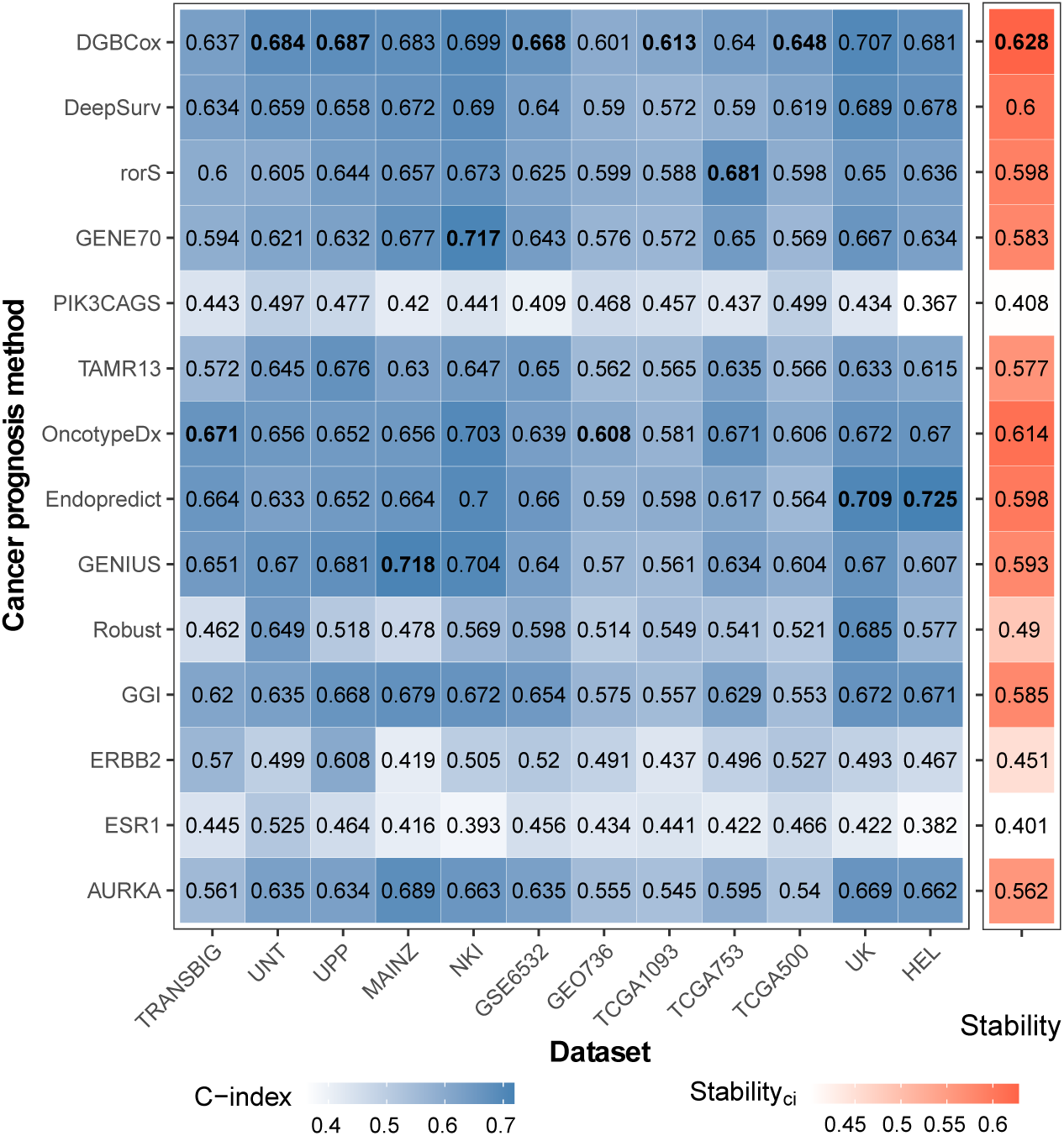
Comparison of different methods for risk score prediction. The left block holds the C-indices of methods on each test dataset. The x-axis stands for the dataset, and the y-axis stands for the method. The C-indices are coloured from white to steel blue. The best result for each data is highlighted in bold. The right block displays *Stability*_*ci*_ of methods across the test datasets. The values of *Stability*_*ci*_ are coloured from white to tomato colour.

When assessing the stability of each method across different environments for the risk score prediction, DGBCox also outperforms all the existing methods because it achieves the highest *Stability*_*ci*_ (0.628) as shown in Fig 4. A good method should have a high average C-index and a low standard deviation of C-indices across test datasets. Conversely, even ESR1 has a low standard deviation (0.038) of C-indices, it performs poorly in most datasets and has a low average C-index (0.439).

We further conduct the comparison of different methods for risk group prediction. As shown in S1 Fig, DGBCox outperforms all the state-of-the-art methods for risk group prediction based on *Stability*_*hr*_ and the number of datasets for which a method has the best *HR*. All the *HR*s of most methods (except ESR1, ERBB2, Robust, and PIK3CAGS) are larger than 1, which means the patients in the predicted high risk group have higher risks of an event when compared to those in the predicted low risk group. These results are consistent with the results of the risk score prediction.

We also use the Log-rank test to evaluate the methods for risk group prediction. The results show that DGBCox successfully stratifies patients in all the twelve test datasets into two risk groups that have distinct survival patterns (S2 Fig). In the meantime, the state-of-the-art methods only succeed on up to nine datasets.

### Important genes for stable prognosis

To further understand which genes are influential in DGBCox, we use the permutation importance algorithm to prioritize genes based on their contributions to prediction stability. Since the signatures discovered by the existing breast cancer prognosis methods (including AURKA, ESR1, ERBB2, GGI, Robust, GENIUS, Endopredict, OncotypeDx, TAMR13, PIK3CAGS, GENE70 and rorS) have been validated as breast cancer related genes, we compare the top 50 ranked genes in DGBCox with these signatures. We map the probe identifiers of signatures to gene symbols if needed. As shown in Fig 5, there are two to three genes in common between the top 50 ranked genes and the breast signature genes identified by Endopredict, OncotypeDx and GENE70 respectively. Four of the 50 genes are confirmed as PIK3CA mutation-associated gene signatures (by PIK3CAGS). Seven of the 50 genes are related to breast cancer histologic grade (by GGI). A significant portion of the top 50 ranked genes of DGBCox is overlapped with the 50 gene signatures of rorS (hypergeometric p-value: 6e-11). Eleven genes are common in the top 50 ranked genes of DGBCox and the 127 genes identified by Robust. Based on the curated list of breast cancer signatures by Huang *et al*. [46], 25 of the top 50 ranked genes have been used for breast cancer prognosis in previous studies.

**Fig 5.**
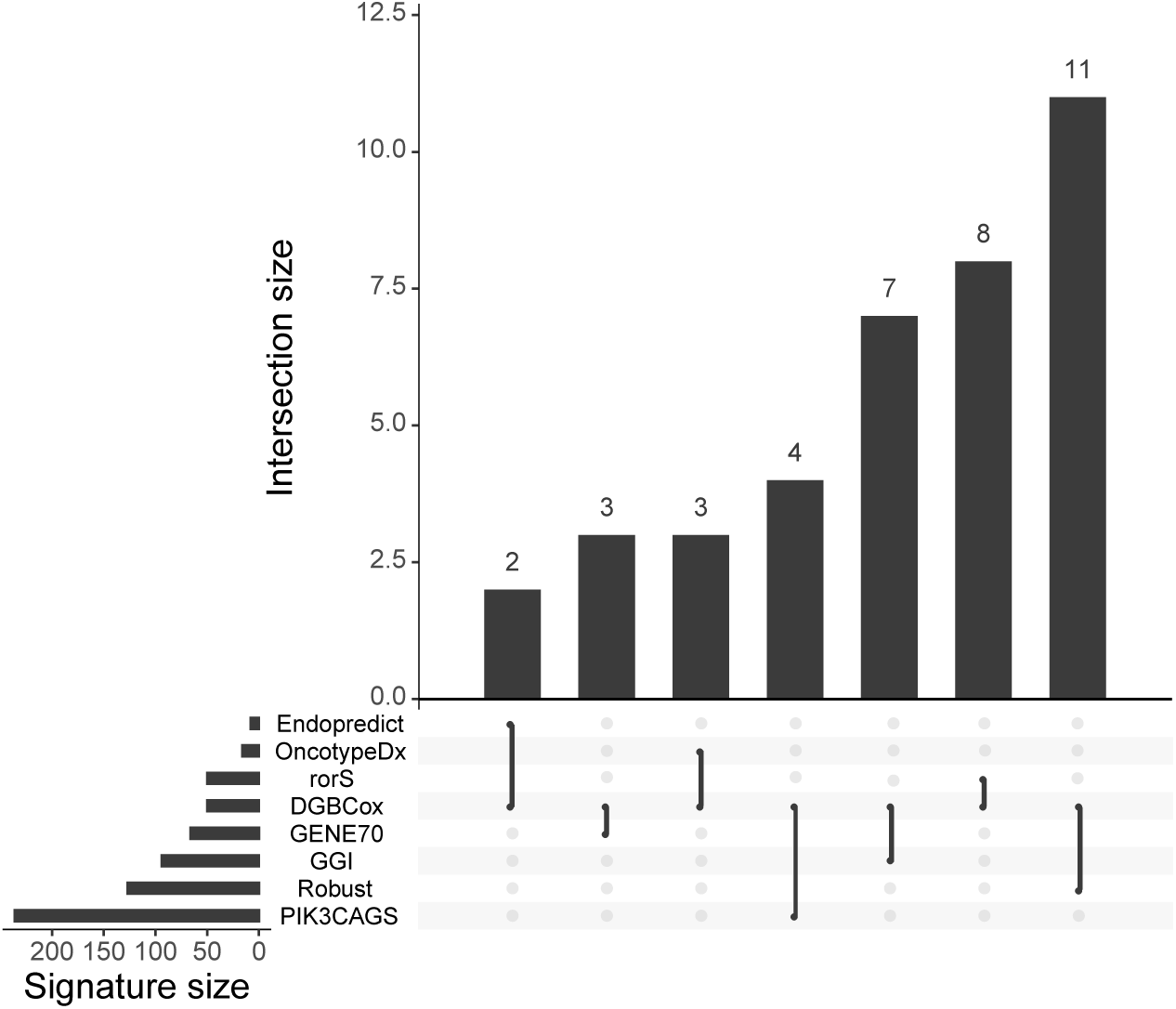
Overlapping genes between DGBCox and the existing methods. The bottom left bar shows the number of signatures in each method. The dotted lines and the diagram on top show that the interaction overlaps between DGBCox and other methods.

We then compare the top 50 ranked genes of DGBCox with cancer causal genes to demonstrate the quality of DGBCox and its findings. It has been confirmed that cancer development and progression are caused by gene mutations [47], and these genes are called cancer causal genes or cancer drivers. We observe that three of the top 50 ranked genes of DGBCox appear in the Cancer Gene Census (CGC) [47], which is a manually curated list of likely cancer drivers. The full list of the top 50 ranked genes and their overlapping with signatures and cancer drivers can be found in S1 Table.

To investigate which biological mechanisms are important for stable breast cancer prognosis, we conduct Gene Ontology (GO) and Kyoto Encyclopedia of Genes and Genomes (KEGG) pathway enrichment analysis (detailed in Section 5 in S1 File). As shown in Fig 6, 16 out of the top 50 ranked genes of DGBCox are mainly enriched in the cell cycle (GO:0140014, GO:0000070 and GO:0007088) and the organelle organization (GO:0000226 and GO:0048285) processes, which are consistent with the functions of most current breast cancer signatures [46]. We also observed some genes are functionally enriched the maternal process involved in female pregnancy (GO:0060135) or the anaphase-promoting complex-dependent catabolic process (GO:0031145). These genes have been reported to be associated with breast cancer development or metastasis. For example, *ESR1* and *PGR* are breast cancer biomarkers that are commonly used for breast cancer subtype diagnosis [48]. *STC2* is reported to be associated with breast cancer cell development [49]. *ANGPT2* is reported to be capable of promoting breast cancer metastasis [50]. *PTTG1* is known as an oncogene that promotes cancer cell proliferation [51], migration, and invasion [52] as well as associated with the epithelial-mesenchymal transition (EMT) in breast cancer [53]. The top 50 ranked genes of DGBCox are significantly enriched (adjusted p-value less than 0.01) in the oocyte meiosis pathway (KEGG ID: hsa04114, as shown in S3 Fig. Several studies have suggested that this pathway plays an important role in the development of breast cancer [54, 55], however, the underlying biological mechanism needs to be verified by further laboratory and clinical researches.

**Fig 6.**
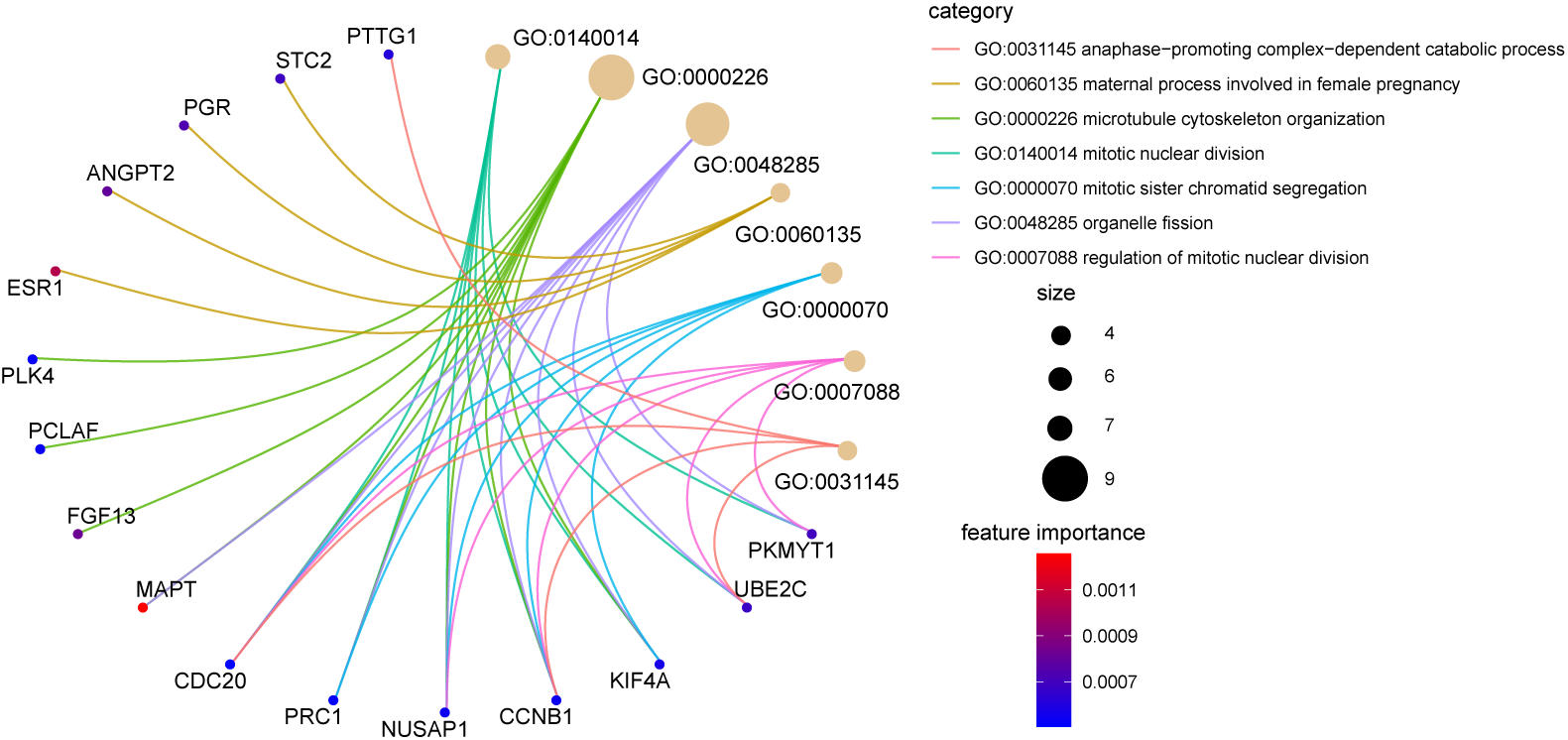
The non-redundant GO terms contained 16 of the top 50 ranked genes. The genes (illustrated by dots) are coloured by their values of permutation feature importance. The size of the circle for a GO term is proportional to the number of genes annotated to this term.

## Conclusion

In this study, we proposed DGBCox to improve the stability of cancer prognosis across unknown environments. To the best of our knowledge, this is the first method designed for the stable cancer prognosis task. Results on 12 independent test datasets demonstrated that DGBCox is effective and stable for breast cancer prognosis across different environments.

We further analyzed which genes are important for the stable breast cancer prognosis. Genes were ranked by the proposed permutation importance algorithm. Interestingly, 25 of the top 50 ranked genes have been used for breast cancer prognosis in previous studies. These 50 genes are significantly enriched in seven non-redundant GO terms and one KEGG pathway which have been reported to be related to the development of breast cancer. The important genes help us understand the molecular mechanism of breast cancer development and also potentially be exploited to improve precision medicine for breast cancer treatment. We believe DGBCox is a useful method in predicting breast cancer prognosis and it will aid researchers in other cancer research fields.

## Supporting information

**S1 File. Supplementary information**.

**S1 Table. The top 50 ranked genes in DGBCox and their overlapping with other studies**. 1: An Entrez ID is a gene’s identifier at the National Center for Biotechnology Information (NCBI)’s database. 2: The gene has been included in Cancer Gene Census (CGC) database [47], which indicates the dysfunction of this gene may cause cancer. 3: The gene has been used for breast cancer prognosis in previous studies based on the list curated by Huang *et al*. [46].

**S1 Fig. Comparison of different methods for risk group prediction**. The left block holds the hazard ratios of methods on each test dataset. The x-axis stands for the dataset, and the y-axis stands for the method. The hazard ratios are coloured from white to steel blue. The best result for each data is highlighted in bold. The right block displays *Stability*_*hr*_ of methods across the test datasets. The x-axis shows the *Stability* _*hr*_, and the y-axis stands for the method. The values of *Stability*_*hr*_ are coloured from white to tomato colour.

**S2 Fig. Comparison of different methods based on the p-values by the Log-rank tests**. The x-axis stands for the dataset, and the y-axis stands for the method. The steel blue tile means that the p-value of a method is less than 0.05, which indicates that the method can successfully stratify patients in the dataset into two risk groups with distinct survival patterns. The grey tile indicates the opposite.

**S3 Fig. The oocyte meiosis pathway (hsa04114) contains six important genes for DGBCox**. These six important genes are highlighted by red colour.

## Acknowledgments

This study makes use of data generated by the Molecular Taxonomy of Breast Cancer International Consortium. Funding for the project was provided by Cancer Research UK and the British Columbia Cancer Agency Branch.

